# Murine polyomavirus microRNAs promote viruria during the acute phase of infection

**DOI:** 10.1101/240994

**Authors:** James M. Burke, Clovis R. Bass, Emin T. Ulug, Christopher S. Sullivan

## Abstract

Polyomaviruses (PyVs) can cause serious disease in immunosuppressed hosts. Several pathogenic PyVs encode microRNAs (miRNAs), small RNAs that regulate gene expression via RNA silencing. Despite recent advances in understanding the activities of PyV miRNAs, the biological functions of PyV miRNAs during *in vivo* infections are mostly unknown. Studies presented here use murine polyomavirus (MuPyV) as a model to assess the roles of the PyV miRNAs in a natural host. This analysis reveals that a MuPyV mutant that is unable to express miRNAs has enhanced viral DNA loads in select tissues at late times after infection, indicating that during infection of a natural host, PyV miRNAs function to reduce viral replication during the persistent phase of infection. Additionally, MuPyV miRNAs promote viruria during the acute phase of infection as evidenced by a defect in shedding during infection with the miRNA mutant virus. The viruria defect of the miRNA mutant virus could be rescued by infecting Rag2-/-mice. These findings implicate miRNA activity in both the persistent and acute phases of infection and suggest a role for MuPyV miRNA in evading the adaptive immune response.

**Importance:** MicroRNAs are expressed by diverse viruses, but for only a few is there any understanding of their *in vivo* function. PyVs can cause serious disease in immunocompromised hosts. Therefore, increased knowledge of how these viruses interact with the immune response is of possible clinical relevance. Here we show a novel activity for a viral miRNA in promoting virus shedding. This work indicates that in addition to any role for the PyV miRNA in long-term persistence, that it also has biological activity during the acute phase. As this mutant phenotype is alleviated by infection of mice lacking an effective adaptive immune response, our work also connects the *in vivo* activity of a PyV miRNA to the immune response. Given that PyV-associated disease is associated with alterations in the immune response, our findings may help to better understand how the balance between PyV and the immune response becomes altered in pathogenic states.

## Introduction

The polyomavirus (PyV) family is comprised of a large number of viruses that predominantly infect vertebrates (1). The founding member of the PyVs, murine polyomavirus (MuPyV), has been the most tractable *in vivo* model for studying the PyV life cycle and has played an important role in understanding PyV-mediated transformation and the antiviral response (2, 3, 4). MuPyV is thought to be transmitted to newborn mice via a respiratory route (5, 6, 7). Primary replication, which occurs 1–6 days post-infection in nonciliated epithelial clara cells of the lungs (7), is followed by dissemination and replication in secondary tissues 7–12 days post infection (6, 8). Viral replication has been observed in various secondary organs, including kidneys, salivary glands, spleen, lymph nodes, heart, liver, skin, lungs, and mammary glands (8, 9, 10). Virus replication peaks around 12 days after infection, concurrent with a rise in viral shedding in the urine and saliva (6). A rise in anti-MuPyV antibody titers occurs 7–15 days p.i. and precedes clearance of detectable virus in most organs by 22–30 days (8).

A reduction in viral loads marks the transition from the acute phase to the persistent phase, which is generally defined by the continued presence of viral DNA in tissues (3). Viral DNA is maintained in select tissues, such as bone marrow, spleen, kidney, salivary glands, and mammary glands (8, 9). The viral DNA in these tissues is thought to represent a persistent reservoir of ‘smoldering’ viral replication in semi-permissive cells (4), although a latent state has yet to be ruled out. A change in the microenvironment (e.g. tissue damage), immune status (e.g. pregnancy or immunosuppression), or hormone levels (e.g. pregnancy, stress), can ‘reactivate’ or increase replication of MuPyV from persistent reservoirs (11, 12). This results in episodic shedding of MuPyV in urine and saliva, which contaminates the surrounding environment with infectious virus (6). Similar observations have been made for human PyVs, whereby increased viruria during pregnancy coincides with a rise in anti-PyV neutralizing antibodies (13, 14, 15).

Many polyomaviruses – including MuPyV, RacPyV, SV40, JCPyV, MCPyV, and BKPyV – encode microRNAs (miRNAs) (16, 17, 18, 19, 20), small regulatory RNAs (~22 nt) that repress gene expression via RNA silencing (21). The PyV miRNAs are encoded in the late orientation, opposite to the T antigen mRNAs. Consequently, miRNAs are perfectly complementary to the T antigen transcripts, permitting Argonaute 2 (Ago2)-mediated cleavage of these mRNAs. This results in a reduction in T antigen protein levels late in infection, a conserved function of PyV miRNAs (16, 17, 18, 19, 20, 22). Despite established roles of T antigen in promoting viral replication, miRNA-null laboratory strains of SV40 and MuPyV replicate at rates similar to their wild type counterparts under standard cell culture conditions (16, 17). However, miRNAs expressed by non-rearranged BKPyV strains, which more closely resemble circulating BKPyV in humans and express lower levels of the early transcripts, are able to downregulate T antigen levels to sufficiently reduce viral replication in primary renal proximal tubule epithelial cells (23). Consistent with these results, MCPyV miRNA promotes long-term persistence by inhibiting DNA replication in a cell culture model (24), and the SV40 miRNAs reduce persistent SV40 DNA loads in Syrian Golden hamster tissues (25). These studies suggest that the PyV miRNAs reduce virus replication by down-regulating T antigen expression. However, the effects of miRNA expression on virus replication in a natural host have yet to be established.

In addition to regulating viral replication, independent studies implicate a role for the PyV miRNAs in inhibiting the host immune response. Downregulation of T-antigen by the SV40 miRNAs decreases CD8 T-cell-mediated lysis in cell culture (16). miRNA-mediated downregulation of the host ULBP3 stress-induced ligand has been suggested to reduce killing of BKV- and JCV-infected cells by natural killer cells (26). MuPyV miRNAs reduce Smad2-mediated apoptosis during infection (27). These studies support a model whereby the PyV miRNAs modulate viral and host transcripts in order to extend the life of infected cells by reducing innate and adaptive immune responses.

Despite advances in understanding the expression and activities of PyV miRNAs, their functions during the viral lifecycle in a natural host are poorly understood. In this study, we use MuPyV as a model to investigate the role of PyV miRNAs on viral dissemination, replication, and shedding during the acute and persistent phases of infection in mice. This work reveals activities of the MuPyV miRNA in both the persistent and acute phases of infection and suggests a link between miRNA function and the adaptive immune response.

## Results

### Sequencing of the MuPyV miRNAs

MuPyV miRNAs were initially identified via secondary structure predictions and confirmed indirectly (17). This approach could only estimate the 5’ and 3’ termini of the derivative miRNAs. It also did not resolve whether MuPyV contains additional miRNA loci. To better characterize the MuPyV miRNAs, we conducted high throughput sequencing of small RNA from NIH3T3 cells infected with the MuPyV PTA strain (Figure 1). These results map with precision the termini of the MuPyV miRNAs and confirm that only the single, previously identified pre-miRNA locus, gives rise to MuPyV-encoded miRNA derivatives.

**Figure 1.**
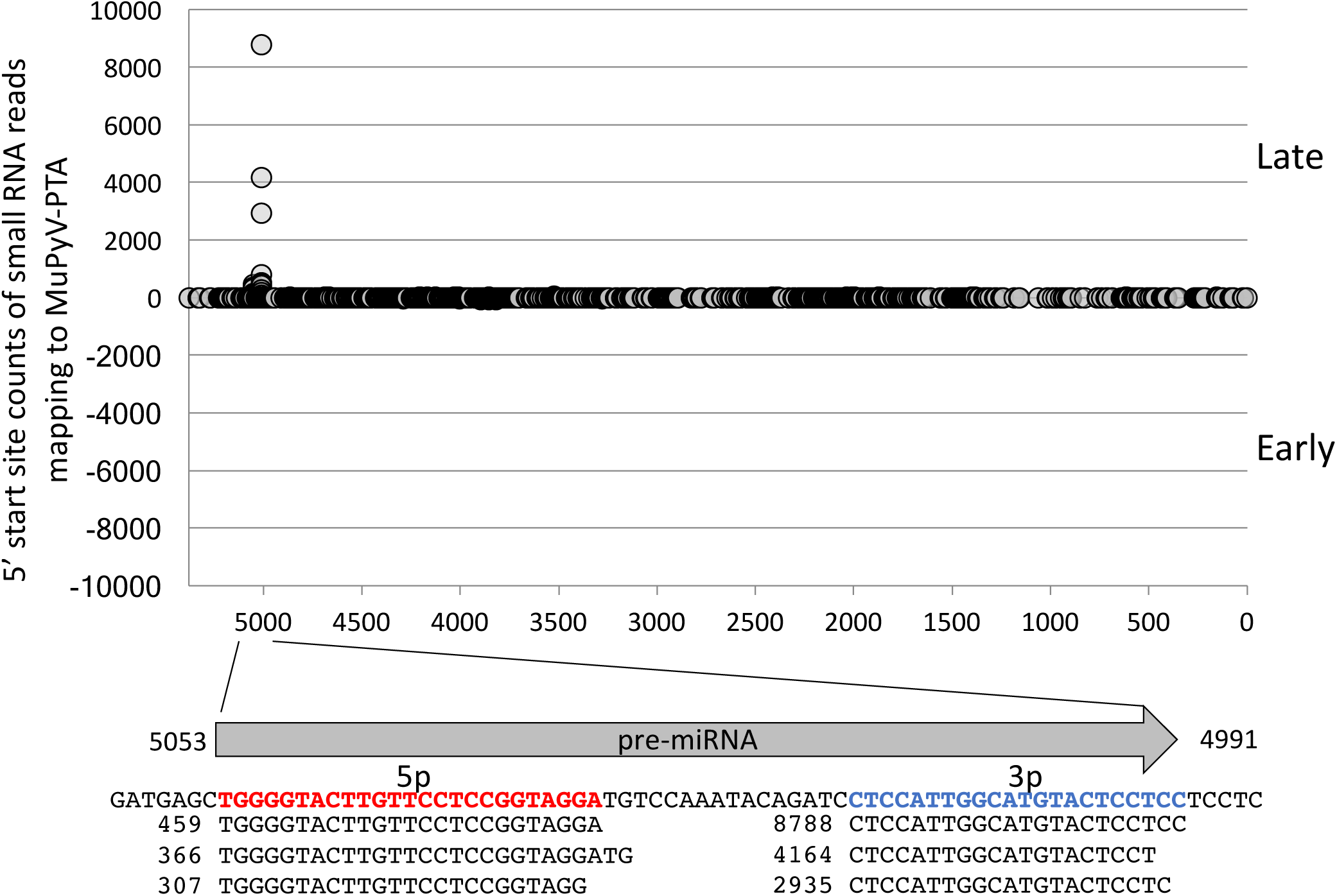
Identification of MuPyV miRNAs by small RNA sequencing. NIH3T3 cells were infected with MuPyV-PTA at an MOI of 0.1. A small RNA library was prepared 7 days p.i. Dots on the graph represent the read counts of the 5’ end of small RNA reads mapping to the MuPyV-PTA genome (NCBI accession number: U27812), positioned on the x-axis. The arrow below the graph represents the pre-miRNA hairpin position and direction. The MuPyV-PTA reference sequence encoding the miRNAs is indicated. Sequences of the three most abundant small RNAs mapping to the 5-prime (bold/red) and 3-prime (bold/blue) arms of pre-miRNA hairpin are shown aligned to the pre-miRNA. Numbers represent the read counts.

### MuPyV miRNAs correlate with reduced viral DNA loads during the persistent phase of infection

PTA-dl1013 is a deletion mutant of the MuPyV-PTA strain that does not express miRNAs (17). Previous studies reported that adult C57BL/6 mice infected with PTA-dl1013 maintained comparable or slightly higher (up to 5-fold) vDNA loads as compared to PTA in spleen and kidney during the acute phase of infection (up to 34 days p.i.) (17). To confirm these results and determine whether the miRNA function at later times post-infection, we examined MuPyV DNA loads by qPCR in spleens, kidneys, and livers of adult C57BL/6 mice infected with 10^5^ IU of PTA or PTA-dl1013 via IP injection during the acute (1 and 4 weeks p.i.) and persistent (10, 16, and 28 weeks p.i.) phases of infection (Figure 2).

**Figure 2.**
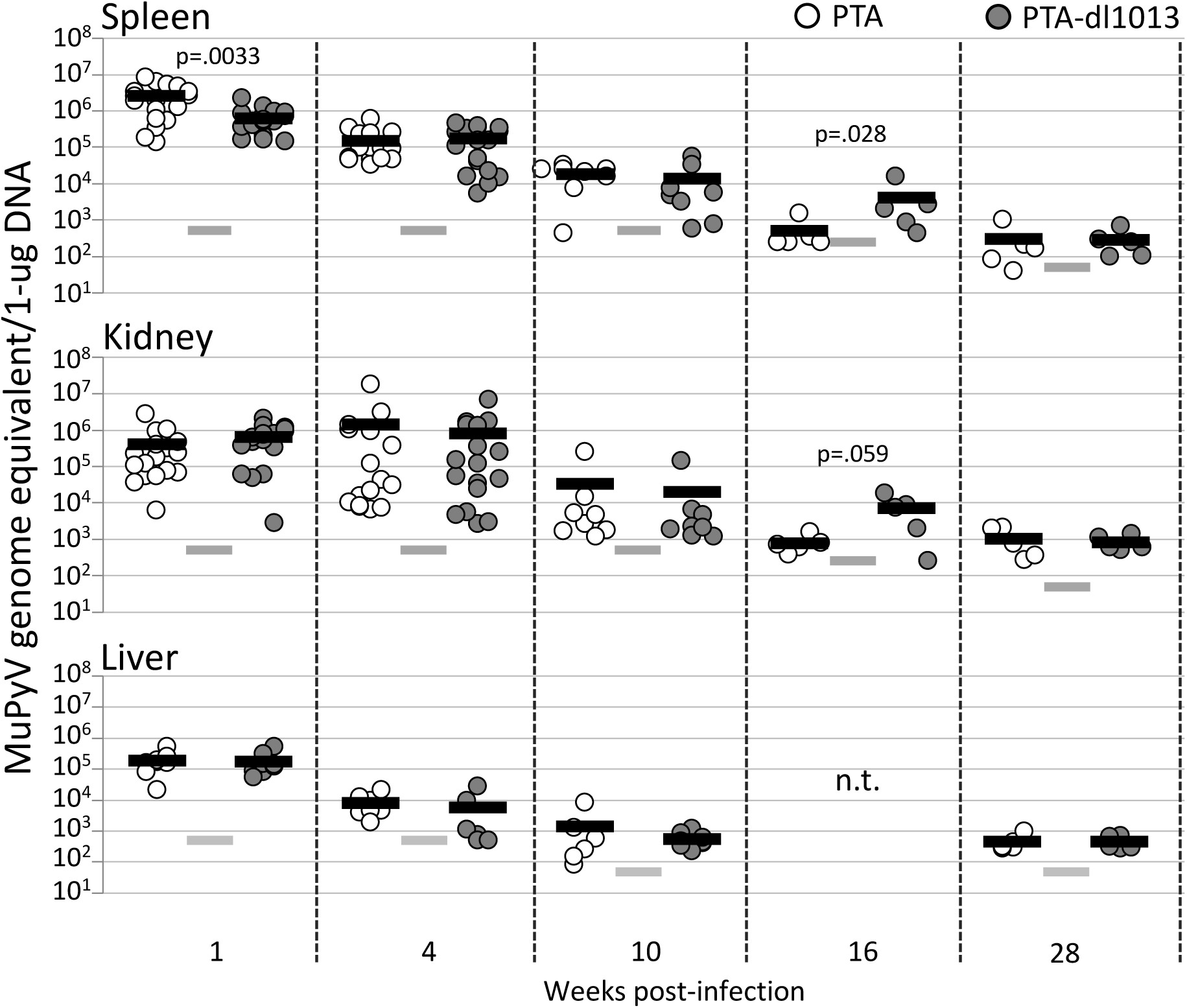
Quantitation of PTA and PTA-dl1013 during the acute and persistent phases of infection. Adult C57BL/6 female mice were inoculated with 1×10^5^ IU of either PTA (white circles) or PTA-dl1013 (gray circles) via IP injection, and genome levels in spleen and kidney were quantified at times post infection by using a TaqMan probe and primers that recognize both genomes. The limit of detection for the qPCR reactions was 10 copies. The input DNA amounts were 20 ng (at 1, 4, and 10 weeks p.i.), 40 ng ( at 16 weeks), or 200 ng (at 28 weeks). Genome copy numbers were normalized per 1ug. Dots represent individual animals. The black bar represents the average genome copy number. The gray bars represent the limit of detection (L.O.D,) of the assay after normalization. Samples that were below the limit of detection are graphed at the limit of detection. P-values were calculated using the Mann-Whitney U test.

High virus yields (2 × 10^5^ −5 × 10^6^ genome equivalents/ug DNA) were observed in spleen and kidney after one and four weeks of infection with PTA, consistent with robust virus replication during the acute phase of replication (Figure 2). Lower levels were observed in liver during these times (Figure 2). Virus loads diminished to ≤ 10^3^ genome equivalents/ug DNA at later times after infection (16 and 28 weeks p.i.), consistent with a low level of virus replication during the persistent phase of replication. These levels were only slightly above the limit of detection of our PCR assay, even when increased input DNA was assayed.

In comparison to PTA, PTA-dl1013 levels were 4.1-fold lower in spleen, but not kidney, at one week p.i. (Figure 2, closed symbols). No significant differences in PTA and PTA-dl1013 DNA levels were detected at 4 weeks or 10 weeks after infection. However, at 16 weeks, PTA-dl1013 genome levels were approximately 10-fold higher than PTA in spleen and kidney. By 28 weeks, only low levels of both PTA and PTA-dl1013 genome levels were detected and no significant differences were observed. These results confirm that the miRNAs are not required for infection and replication in tissues during the acute phase of infection, and indicate that miRNA expression correlates with reduced genomic DNA in certain tissues during the persistent phase of infection.

The results summarized in Figure 2 reveal a substantial variation in viral copies between different mice. To minimize variations due to inter-host differences, we compared levels of PTA and PTA-dl1013 in individual animals. C57BL/6 mice were co-infected with equivalent amounts of PTA and PTA-dl1013 by IP injection. Tissues were then harvested during the acute (2 and 4 weeks) and persistent (10 and 16 weeks) phases of infection, and genome levels were determined by strain-specific qPCR (Supplemental Figure 1A-C). Similar to individually infected mice, we observed high levels of both PTA and PTA-dl1013 in tissues during the acute phase of infection, with MuPyV levels decreasing during the persistent phase of infection (Figure 3). We note that PTA was not detected in the spleen or kidney by 16 weeks p.i., while PTA-dl1013 was present at 10^3^ – 10^4^ copies/ug DNA (Figure 3). To compensate for variation in different animals, we calculated the ratio of PTA to PTA-dl1013 genomes in individual mice. This analysis reveals that during the acute phase of infection (2 and 4 weeks p.i.), the levels of PTA and PTA-dl1013 were comparable in spleen and kidney (Figure 4). At later times, PTA-dl1013 copy numbers were 10-fold higher than PTA in the spleen and kidney at weeks 10 and 16 and 100-fold higher in the kidney at week 16 (Figure 4). We did not observe substantial differences in the liver, but did observe 7-fold higher PTA-dl1013 in the bladder at week 4 (Figure 4). These results are consistent with the elevated levels of PTA-dl1013 detected in the spleen and kidney at 16 weeks following infections with the individual virus strains (Figure 2) and suggest that MuPyV miRNA expression correlates with reduced viral loads in select tissues during the persistent phase of infection.

**Figure 3.**
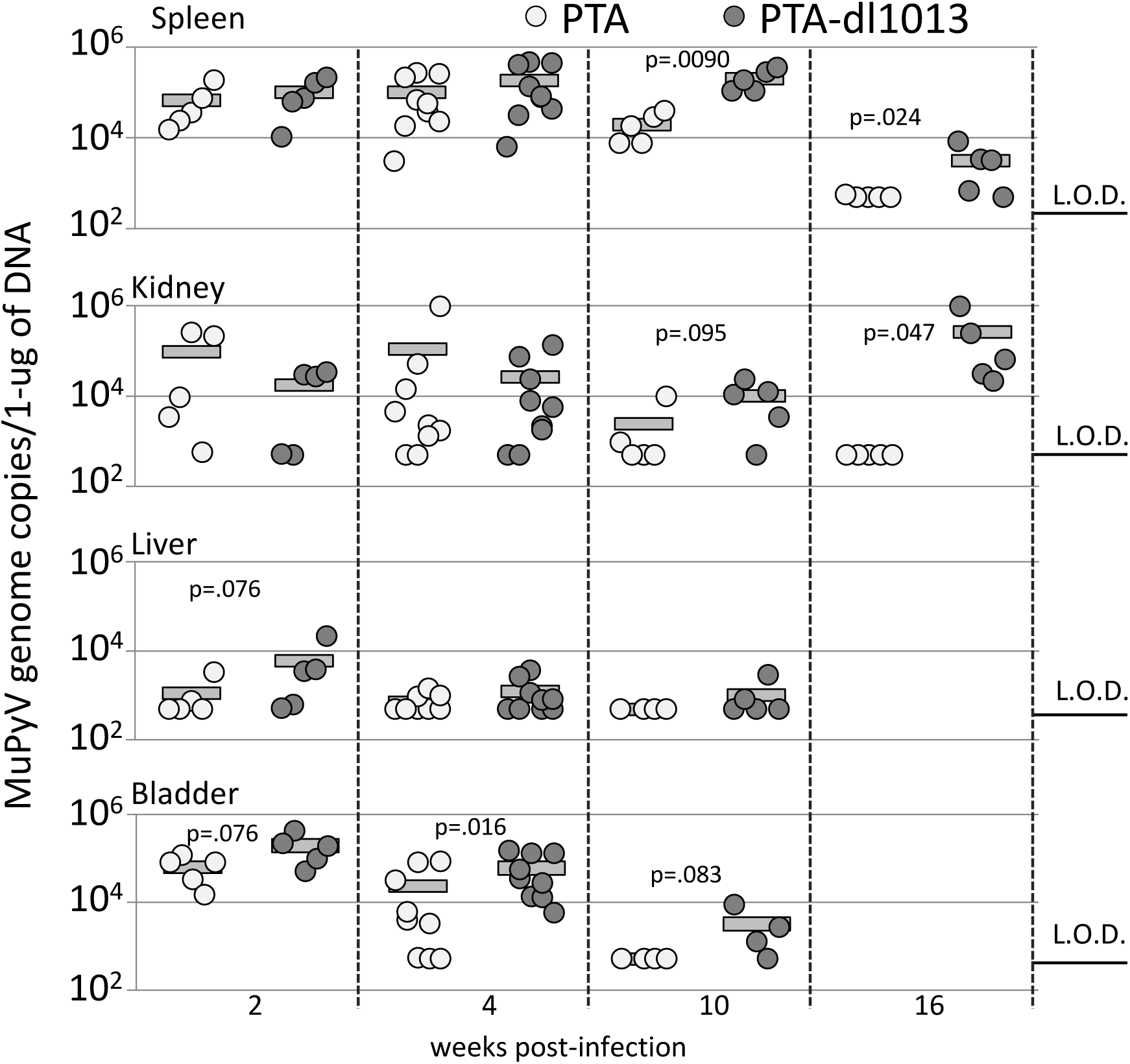
Quantitation of PTA and PTA-dl1013 in tissues of co-infected mice. qPCR analysis of PTA and PTA-dl1013 genome levels in the spleen, kidney, liver, and bladder of adult C57BL/6 mice inoculated with 5×10^4^ IUs of PTA and 5×10^4^ IUs of PTA-dl1013 via IP injection. Tissues were harvested at indicated time points. DNA was recovered and subject to qPCR analysis using PTA- and PTA-dl1013-specific probes. The limit of detection (L.O.D.) was 10 copies. Samples in which PTA or PTA-dl1013 were below the limit of detection were graphed at the limit of detection. The quantities were normalized to 1-ug of total DNA. P-values were determined using Mann-Whitney U test.

**Figure 4.**
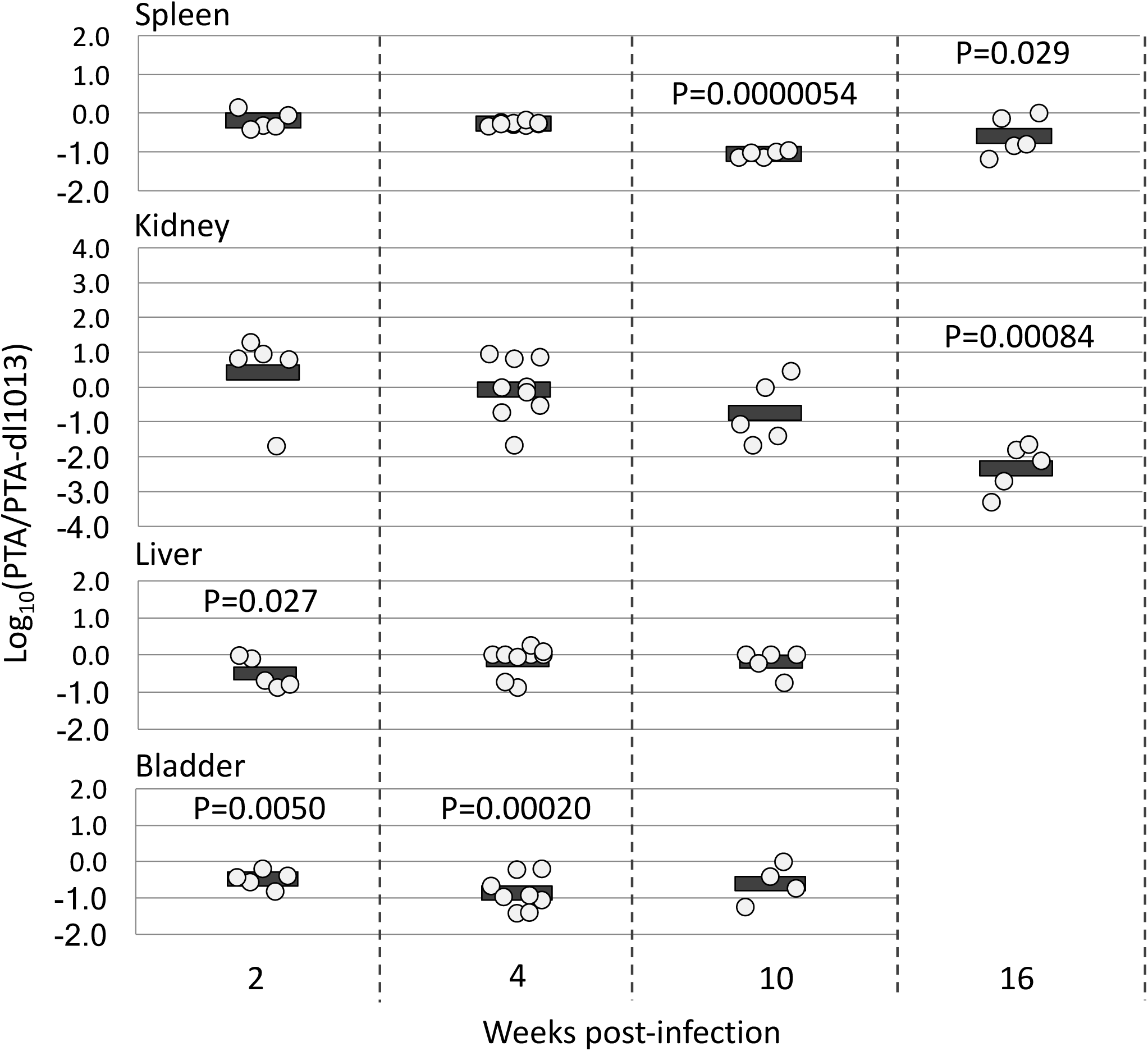
Ratio of PTA and PTA-dl1013 in tissues of co-infected mice. The ratio of PTA and PTA-dl1013 genome copies from (A) in the indicated organs of individual mice co-infected with PTA and PTA-dl1013. Values below the limit of detection were set at the limit of detection. The dots represent Log_10_(PTA/PTA-dl1013). P-values were calculated using one-sample t-test.

### MuPyV miRNAs promote viruria during the acute phase of infection

Early epidemiological studies of MuPyV revealed that high titers of infectious virus are excreted in the urine of mice during the acute phase of infection and then episodically shed during the persistent phase of infection (6). The periodic shedding of MuPyV and many other PyVs in the urine is considered a primary route of transmission (6,28). Therefore, we investigated whether the MuPyV miRNAs affects the magnitude and/or frequency of MuPyV shedding in the urine during the acute and persistent phases of infection. To test this, we collected urine specimens periodically post-infection and quantitated viral genomes. To assess whether PTA-dl1013 could maintain long-term persistence, we treated mice with cyclosporine at day 474 post-infection in order to immunosuppress the mice, which is correlated with increased PyV replication (4).

In the majority of mice individually infected with PTA or PTA-dl1013, we observed vDNA shedding of both strains during the acute phase of infection (13–34 days p.i.) (Supplemental Figure 2), indicating that the MuPyV miRNAs are not required for establishing viruria. Both strains were episodically shed throughout the persistent phase (34–87 day p.i.), which demonstrates that the MuPyV miRNAs are not required for persistent viruria (Supplemental Figure 2). The magnitude and timing of virus shedding by individual mice infected varied substantially (Supplemental Figure 2), and thus no statistical differences between PTA and PTA-dl1013 levels could be determined.

To circumvent the differences in shedding due to variations between individual mice, we tested whether expression of miRNA could correlate with viral shedding in mice co-infected with both PTA and PTA-dl1013. Adult C57BL/6 female mice were co-infected with equal amounts of the PTA and PTA-dl1013, and PTA and PTA-dl1013 DNA was quantitated during the acute and persistent phases of infection (Figure 5A). We did not detect substantial quantities of either PTA or PTA-dl1013 genomes in the urine of ~40% of the mice at any time point during the acute phase of infection (Supplemental file). The remaining “high-shedder” mice expressed readily detectable MuPyV DNA in the urine. MuPyV in these samples was detected at high levels beginning by 10 days p.i. and peaked between 13–27 days, after which the number of shedding mice and the magnitude of shedding began to decrease. Thereafter, we observed episodic shedding events throughout the persistent phase of infection (34–87 days p.i.). These findings reveal that while some mice rarely or never shed detectable virus under our assay conditions, ~60% that are competent for acute and persistent viruria.

**Figure 5.**
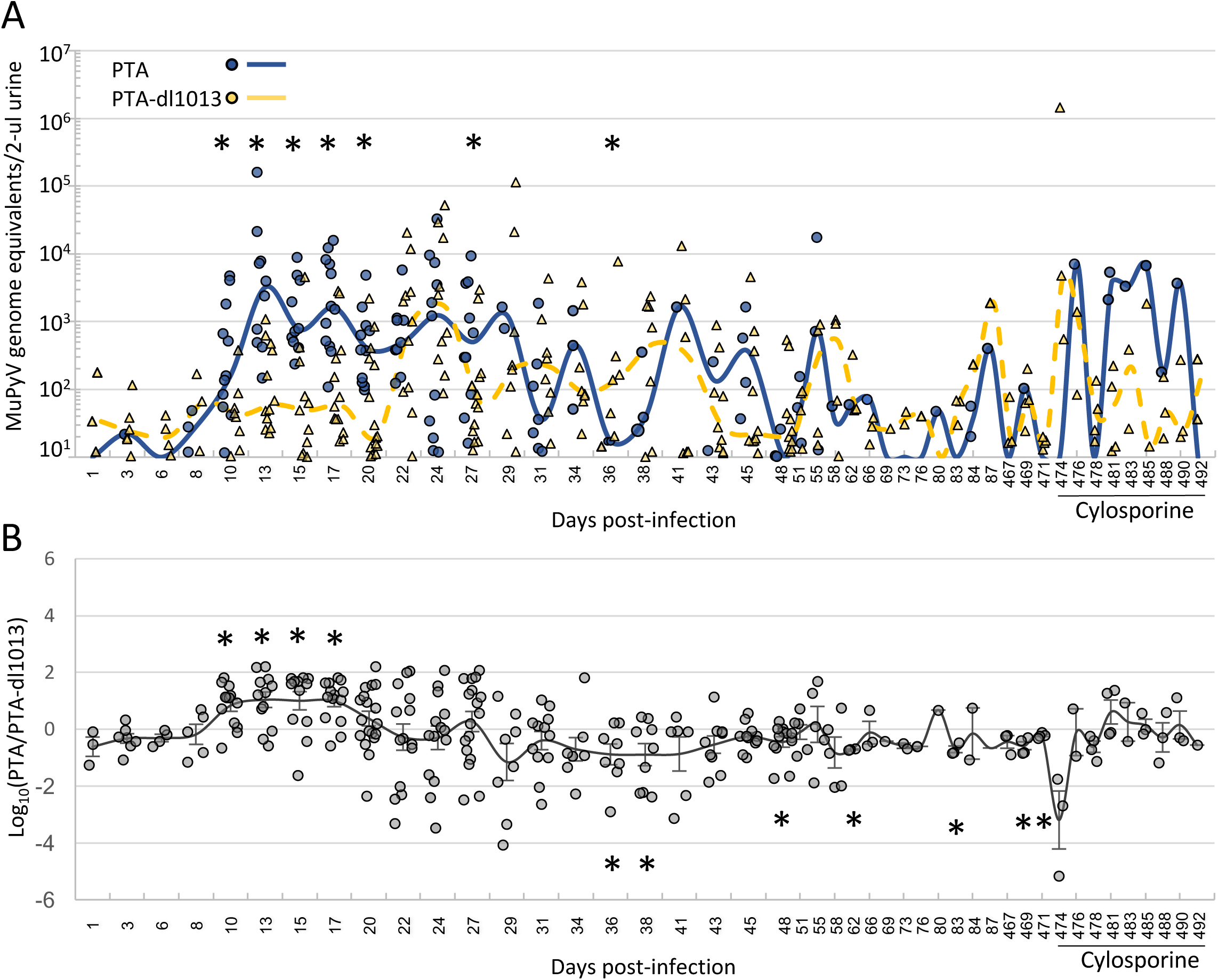
Quantitation of PTA and PTA-dl1013 DNA shedding in the urine. Mice were co-infected with 5×10^4^ IU of PTA and 5×10^4^ IU of PTA-dl1013 via IP injection, urine samples were collected at the indicated times, and strain-specific qPCR was used to quantify wt and mutant genomes. The number of mice tested at each time ranged from 18–30 mice during days 1–27 p.i., 10–20 mice during days 29–55 p.i., and 4 mice during days 62–492 p.i. Panel A presents a dot plot of genome equivalents of PTA (blue circles) and PTA-dl1013 (yellow triangles) in 2 ul of urine from mice that shed viral genome copies equal to or greater that the limit of detection of the qPCR assay. Mice that did not shed above the limit of detection or time points in which genomes were not detected were not included in this analysis. Lines represent the median values of PTA (solid blue) and PTA-dl1013) (dotted yellow). Significant differences (P ≤ 0.05) between PTA and PTA-dl1013 genome levels are marked by asterisks. Panel B compares the ratio of wt and mutant genomes in individual mice co-infected with PTA and PTA-dl1013. When only one genome could be detected (either PTA or PTA-dl1013), the other genome was set at the limit of detection in order to calculate the PTA/PTA-dl1013 ratio. The line represents the average Log_10_(PTA/PTA-dl1013) ± SEM. P values were calculated by using a one sample t-test (asterisks indicate P ≤ 0.05).

The median PTA levels were significantly higher (5- to 100- fold) than PTA-dl1013 during the majority of the acute phase of infection (10, 13. 15, 17, 20 and 27 days p.i.) in co-infected mice (Figure 5A). To minimize the variance in individual mice, we determined the ratio of PTA:PTA-dl1013 DNA in each individual mouse. This data confirmed that PTA was shed at significantly higher levels on average (~10-fold) than PTA-dl1013 during days 10–17 of the acute phase of infection in individual mice co-infected mice (Figure 5B). We did not observe statistically significant differences in the magnitude of viral shedding at the majority of times tested during the persistent phase of infection (Figure 5A, 5B, Supplemental file). Although this may be due to high variance in the timing and the magnitude of shedding, as well as the lower number of mice tested, this suggests that the MuPyV miRNAs do not promote viruria during the persistent phase of infection. In fact, we observed eighty-four PTA-dl1013 shedding events that were above the limit of detection from days 34 through 87, compared to thirty-three for PTA (Figure 5A). This implies that the MuPyV miRNAs may limit detectable shedding during persistent phase of infection, consistent with the reduced PTA levels in tissues (Figures 2–4). At late times post-infection (467-492), we observed PTA and PTA-dl1013 shedding events in a small number of mice. Thus, the MuPyV miRNAs are not required to maintain long-term persistent infections. These data further indicate that the MuPyV miRNAs promote viral shedding during the acute phase of infection, but limit the number of MuPyV shedding event during the persistent phase of infection.

### The shedding defect of miRNA-null MuPyV is mitigated in Rag2-/-mice

The host immune response plays an important role in controlling MuPyV infection (2–4), and two lines of evidence suggest that PyV miRNAs may function to reduce the adaptive immune response. First, miRNA-mediated down-regulation of T-antigens decreases susceptibility to cytotoxic T-cell-mediated lysis to SV40-infected cultured cells (16). Second, miRNA-null strains of SV40 and JCV have only been isolated from severely immunocompromised hosts, suggesting that an active immune response provides selective pressure to maintain miRNA expression (29). Since the absence of the MuPyV miRNA results in decreased virus shedding during the acute phase of infection, we tested whether the defect in acute viral shedding of PTA-dl1013 is mitigated in C57BL/6 *Rag2-/-* mice, which lack mature T- and B-cells. C57BL/6 *Rag2-/-* mice were co-infected with PTA and PTA-dl1013, and the shedding of DNA in urine was monitored. Unlike wild type mice, where ~40% of animals did not shed detectable vDNA during the acute phase of infection, all infected C57BL/6 *Rag2-/-* mice shed detectable levels of MuPyV by 10 days postinfection (Figure 6 and Supplemental file). Levels of both PTA and PTA-dl1013 DNA in urine were found to continually increase in the co-infected C57BL/6 *Rag2-/-* mice up to 20 days p.i., whereas MuPyV levels leveled off and began to decrease by 20 days p.i. (Figure 6). These observations are consistent with the host adaptive immune response being a major factor that controls MuPyV shedding. Importantly, unlike in wild type C57BL/6 mice, C57BL/6 *Rag2-/-* mice shed PTA-dl1013 DNA at levels similar to PTA during the acute phase of infection (10–13 days p.i.) (Figure 6). These data indicate that the MuPyV miRNAs promote viruria during the acute phase of infection in a manner that is dependent on the host having an intact adaptive immune response.

**Figure 6.**
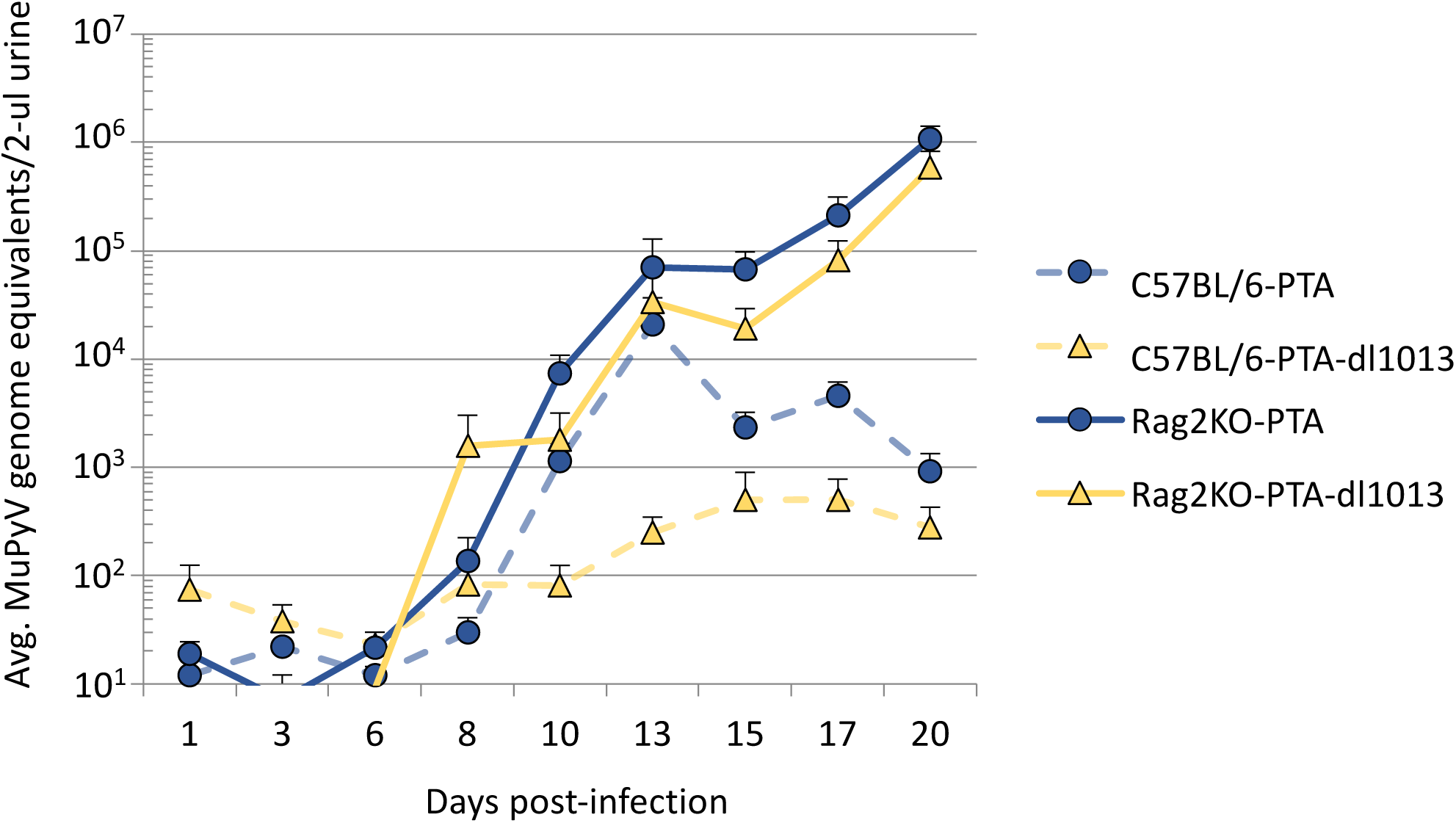
Analysis of PTA and PTA-dl1013 shedding in C57BLC57bl/6 *Rag2-/-* mice. Nine female C57BL/6-RAG2-/- mice were co-infected with 5 ×10^4^ IU of PTA and 5 ×10^4^ IU of PTA-dl1013 via IP injection. Urine was collected during the acute phase of infection, and genomic DNA was quantified by strain-specific qPCR analysis. Wild type C57BL/6 mice co-infected with PTA (blue circles/dashed line) and PTA-dl1013 virus (yellow triangles/dashed line) are from Figure 6. PTA genomes (blue circles/solid line) and PTA-dl1013 virus (yellow triangles/solid line) from C57BL/6-Rag2-/- mice are presented. Plots represent the average (±SEM, n=8–9) at the indicated times.

## Discussion

The natural *in vivo* functions of PyV miRNAs are poorly understood. Results presented here reveal that, consistent with previous studies in cell culture models and a non-natural host (23–25), the PyV miRNAs function to reduce viral loads during the persistent infection of a natural host. In addition, we uncover novel effects of the MuPyV miRNAs on viruria during the persistent and acute phases of infection that provide new insights into PyV miRNA biology.

Similar to previous studies (17), we did not observe large differences in the gross levels of PTA (wild type) and PTA-dl1013 (miRNA-null) strains of MuPyV in tissues during acute phase of infection (weeks 1, 2 and 4 p.i.). This further confirms that the MuPyV miRNAs are not required for acute viral infection and replication *in vivo* under laboratory conditions. However, at weeks 10 and 16 p.i., we observed higher levels of PTA-dl1013 DNA, relative to PTA, in kidney and spleen of mice following single- or co-infection with PTA and PTA-dl1013 (Figures 2–4). Similar findings were observed in tissues of Syrian golden hamsters infected with miRNA-null strains of the simian virus SV40 (25). These results are consistent with the notion that miRNA-mediated downregulation of T antigen reduces virus replication in specific tissues/cell-types at late times after infection to promote viral persistence (23,24).

Our assessment of MuPyV viruria revealed that the MuPyV miRNAs are not required to establish or maintain long-term viral persistence (Figure 5 and Supplemental Figure 2). However, we did observe more PTA-dl1013 than PTA shedding events during the persistent phase of infection (34-87 days p.i.). This is consistent with increased PTA-dl1013 levels in tissues and further indicates that the MuPyV miRNAs may limit viral replication events during the persistent phase of infection. Though the function of this activity remains unclear, it may be important to prevent priming of the host immune response in order to transmit to naïve hosts more efficiently. Alternatively, this activity could prevent viral-associated pathologies to the host, which would presumably decrease viral fitness. Future experiments will address the relevant role of miRNA-mediated reduction of viral loads in urine and tissues during the persistent phase of infection.

During the acute phase of infection (days 10–20) in wild type C57BL/6 mice, only ~60% of mice shed substantial levels of MuPyV DNA (Figure 5 and Supplemental Figure 2). This is in contrast to infections in immunodeficient C57BL/6 *Rag2-/-* mice, in which 100% of mice shed substantial levels of MuPyV DNA. This suggests that the adaptive immune response is a major factor controlling viruria. Analysis of wild-type C57BL/6 mice co-infected with PTA and PTA-dl1013 revealed that PTA-dl1013 DNA was shed at significantly lower levels than PTA (Figure 5), which indicates the MuPyV miRNAs promote viral shedding at early times post-infection. Importantly, the defect in PTA-dl1013 was largely mitigated during infection of immunodeficient C57BL/6 *Rag2-/-* mice (Figure 6), suggesting that the MuPyV miRNAs help evade a component of the host adaptive immune response to promote viruria. We note that PTA-dl1013 shedding was not fully rescued by infecting C57BL/6 *Rag2-/-* mice, possibly indicating that the MuPyV miRNAs have additional activities [for example, reduction of smad2-mediated apoptosis (27)] in promoting viruria independent of the adaptive immune response. These combined data demonstrate that the MuPyV miRNAs promote viral shedding by at least indirectly interacting with the adaptive immune response. This activity could be important in the wild as a means to increase transmission of the virus to susceptible hosts during the acute phase of infection.

The exact mechanism(s) by which MuPyV miRNA expression enhances virus shedding in immunocompetent mice remains unclear. Because we observed miRNA-dependent promotion of shedding in co-infected immunocompetent mice, this indicates that MuPyV miRNAs are likely functioning via a cell autonomous mechanism to repress the effects of the adaptive immune response. This is consistent with previous findings that the SV40 miRNAs can reduce CD8 T-cell-mediated lysis. However, it is unclear why the miRNAs promote viruria only during a narrow window of time post-infection. One possibility is that the MuPyV miRNAs increase dissemination to the target cells required for subsequent viruria, possibly by reducing the CTL response to promote infection/replication in lymphocytes early during the acute phase of infection. In support of this, at one week p.i., but not at 2 and 4 weeks p.i., we observed ~4-fold higher PTA DNA levels specifically in the spleen (Figures 2–4), indicating the MuPyV miRNAs may promote infection of lymphocytes early during infection. Alternatively, the cell-mediated adaptive immune response to MuPyV may change over time, making the miRNAs less effective at promoting viruria. While the CD8+ T cell response to MuPyV is predominately to T-antigen, the CD8+ T cell response has also been shown to be directed toward a VP2–derived nonamer peptide (30). Thus, as the infection progresses, miRNA-mediated down-regulate of T-antigens may become less effective at muting the CTL response as the host T-cell response develops more so against the late VP proteins. Future studies are required to more precisely define the relevant components of the immune response important for hindering the miRNA mutant virus.

In summary, this study demonstrates functionality of MuPyV miRNAs during both the persistent and acute phases of infection and at least indirectly implicates their function in altering some component of the adaptive immune response. Given the clinical relevance of understanding the interplay between polyomaviruses and the immune response, this work provides a useful, quantitative, and non-invasive system for future efforts at understanding PyV miRNA effects on adaptive immunity.

## Materials and Methods

### Cell cultures and virus strains

Primary baby mouse kidney (BMK) cell cultures were prepared from 2–4 week old female C57BL/6 mice (Jackson Laboratories). Kidneys were excised, washed in PBS, and incubated in DMEM containing 0.25% trypsin overnight at 4°C. Kidneys were homogenized, incubated in trypsin at 37°C with agitation, and suspended in DMEM+10% FBS (Cellgro). Cells were washed by centrifugation and plated at a density of 10^7^ cells per 100 mm dish. NIH3T3 cells were maintained in DMEM+10% FBS.

The PTA and PTA-dl1013 virus strains have been described previously (17). Virus was propagated by infecting primary BMK cells (80% confluent) with PTA or PTA-dl1013 at an MOI of 0.05. Virus stocks were prepared after 10 days by scraping the cells into a portion of the culture medium. The cell extracts were then subjected to three freeze/thaw cycles, clarified by centrifugation, and stored at −80°C.

Virus titers were determined by immunofluorescence. 80–90% confluent NIH3T3 cells in 12- or 24-well plates were infected with dilutions of the PTA and PTA-dl1013 virus stocks for one hour. Infected cells were maintained in DMEM+2% FBS for 40 hours and then fixed and permeabilized with 4% paraformaldehyde and 1% NP-40. After blocking with 10% goat serum overnight, cells were incubated with rabbit-anti-PyV antibody (a gift from Dr. Richard Consigi) for 30 min. After three PBS washes, cells were stained with Cy3-goat-anti-rabbit IgG (Abcam) for 10 min. and washed. The numbers of total and stained cells per field were quantified by using an inverted fluorescent microscope (Leica), and infectious titers (IU/ml) were determined.

### Animal infections and tissue analysis

Female C57BL/6 (Jackson Laboratories) and C57BL/6 *Rag2-/-* mice were inoculated with 1 × 10^5^ IU of either PTA or PTA-dl1013 via intraperitoneal (IP) injection between 4 and 5 weeks of age. For co-infections, mice were inoculated (IP) with 5 × 10^4^ IU of PTA and 5 × 104 IU of PTA-dl1013. Cylcospoin was administered as described in (31). Tissues were harvested at the indicated times and flash frozen in liquid nitrogen. Homogenates were prepared by grinding with a mortar and pestle chilled with liquid nitrogen. DNA was extracted from 25 mg of tissue by using the DNeasy Blood and Tissue kit (Qiagen). Recovered DNA was quantitated by the Nanodrop procedure and diluted to 100 ng/ul or 10 ng/ul for qPCR analysis.

### Urine analysis

To collect urine, mice were placed on wire-bottomed cages for 2–4 hours. Urine was collected on Parafilm, and 50 ul aliquots (or the total if less than 50 ul) were used for DNA extraction with the QIAamp viral RNA mini kit (Qiagen) following the manufacturer’s protocol to remove PCR inhibitors. DNA was eluted in 60 ul of TE buffer. MuPyV copy numbers were normalized to that in 2 ul of urine.

### Quantitative PCR

Concentrated pBluescript-sk+PTA and pBluescript-sk+PTA-dl1013 vectors were obtained via Maxi Prep (Invitrogen). Plasmids were prepared at 85.5 ng/ul (10^10^ copies of PTA or PTA-dl1013 per ul) and then serial diluted. For tissue analysis from individually infected mice, 20 ng of DNA (for samples taken 1, 4, 10 weeks p.i.) or 200 ng of DNA (10 and 28 weeks p.i.) in 2 ul of TE buffer was added to an 8 ul reaction mixture containing 5 ul of 2X gene expression master mix (Applied Biosystems), 0.75 uM of MuPyV sense primer (CGCACATACTGCTGGAAGAAGA), 1.0 uM MuPyV antisense primer (TCTTGGTCGCTTTCTGGATACAG), and 100 nM of MuPyV TaqMan MGB (Applied Biosystems) probe (FAM-ATCCTTGTGTTGCTGAGCCCGATGANFQ) as described (32). For specific detection of PTA and PTA-dl1013 in urine, 2 ul of DNA (of the 60 ul extracted unless otherwise indicated in Supplemental file) was added to an 8 ul reaction mixture containing 5 ul 2X gene expression master mix (Applied Biosystems), 0.75 uM of PTA/PTA-dl1013 sense primer (GATGAGCTGGGGTACTTGT), 0.75 uM of PTA/PTA-dl1013 antisense primer (TGTATCCAGAAAGCGACCAAG), and 100 nM of either the PTA-specific TaqMan MGB (Applied Biosystems) probe (FAM-TAGGATGTCCAAATACAGATCCTCNFQ) or PTA-dl1013-specific TaqMan MGB (Applied Biosystems) probe (FAM-CTCCGGTTCCATTGGCATGT-NFQ). The PTA and PTAdl1013 standards (107/ul -101/ul) were confirmed by using the universal MuPyV primers/probe, which detects both PTA and PTA-dl1013, to ensure that the copy number of each standard was equal. Assays were performed using 384-well format plates in a ViiA™ 7 Real-Time PCR System (Applied Biosystems). The limit of detection was 10 copies.

### Statistical analyses

One sample T-test (two-tailed) and Mann-Whitney U-test (two-tailed) were performed using real statistics resource package for excel (http://www.real-statistics.com/).

## Acknowledgements

The work described in this article was supported by a Burroughs Wellcome Investigators in Pathogenesis Award to CSS and a grant from the Cancer Prevention and Research Institute of Texas [RP140842]. The authors would like to thank Dr. Lauren Ehrlich and Dr. Edward Marcotte for reagents, and Dr. Luis Villarreal for insights regarding viral persistence.

## References

1. Johne R, Buck CB, Allander T, Atwood WJ, Garcea RL, Imperiale MJ, Major EO, Ramqvist T, Norkin LC. 2011. Taxonomical developments in the family Polyomaviridae. Arch Virol 156:1627–1634.

2. Benjamin TL. 2001. Polyoma virus: old findings and new challenges. Virology 289: 167–173.

3. Ramqvist T, Dalianis T. 2009. Murine polyomavirus tumour specific transplantation antigens and viral persistence in relation to the immune response, and tumour development. Semin Cancer Biol 19: 236–243.

4. Swanson PA, Lukacher AE, Szomolanyi-Tsuda E. 2009. Immunity to polyomavirus infection: The polyomavirus–mouse model. Semin Cancer Biol 19: 244–251.

5. Rowe WP, Hartley JW, Brodsky I, Huebner RJ, Law LW. 1958. Observations on the spread of mouse polyoma virus infection. Nature 182:1617–16179.

6. Rowe W P. 1961. The epidemiology of mouse polyoma virus infection. Bacteriol Rev 25:18–31.

7. Gottlieb K, Villarreal LP. 2000. The Distribution and Kinetics of Polyomavirus in Lungs of Intranasally Infected Newborn Mice. Virology 266: 52–65.

8. Dubensky TW, Villarreal LP. 1984. The primary site of replication alters the eventual site of persistent infection by polyomavirus in mice. J Virol 50: 541–546.

9. Wirth JJ, Amalfitano A, Gross R, Oldstone MB, Fluck MM. 1992. Organ- and age-specific replication of polyomavirus in mice. J Virol 66:3278–86.

10. Berke Z, Dalianis T. 1993. Persistence of polyomavirus in mice infected as adults differs from that observed in mice infected as newborns. J Virol 67: 4369–4371.

11. Atencio IA, Shadan FF, Zhou XJ, Vaziri ND, Villarreal LP. 1993. Adult mouse kidneys become permissive to acute polyomavirus infection and reactivate persistent infections in response to cellular damage and regeneration. J Virol 67: 1424–1432.

12. McCance DJ, Mims CA. 1979. Reactivation of polyoma virus in kidneys of persistently infected mice during pregnancy. Infect Immun 25: 998–1002.

13. Coleman DV, Gardner SD, Mulholland C, Fridiksdottir V, Porter AA, Lilford R, Valdimarsson H. 1983. Human polyomavirus in pregnancy. A model for the study of defense mechanisms to virus reactivation. Clin Exp Immunol 53: 289–296.

14. Eash S, Manley K, Gasparovic M, Querbes W, Atwood WJ. 2006. The human polyomaviruses. Cell Mol Life Sci 63:865–876.

15. Gibson PE, Field AM, Gardner SD, Coleman DV. 1981. Occurrence of IgM antibodies against BK and JC polyomaviruses during pregnancy. J Clin Pathol 34: 674–679.

16. Sullivan CS, Grundhoff AT, Tevethia S, Pipas JM, Ganem D. 2005. SV40-encoded microRNAs regulate viral gene expression and reduce susceptibility to cytotoxic T cells. Nature 435: 682–686.

17. Sullivan CS, Sung CK, Pack CD, Grundhoff AT, Lukacher AE, Benjamin T, Ganem D. 2009. Murine polyomavirus encodes a microRNA that cleaves early RNA transcripts but is not essential for experimental infection. Virology 387: 157–167.

18. Seo GJ, Chen CJ, Sullivan CS. 2009. Merkel cell polyomavirus encodes a microRNA with the ability to autoregulate viral gene expression. Virology 383: 183–187.

19. Seo GJ, Fink LH, O’Hara B, Atwood WJ, Sullivan CS. 2008. Evolutionarily conserved function of a viral microRNA. J Virol 82: 9823–9828.

20. Chen CJ, Cox JE, Azarm K, Wylie KN, Woolard KD, Pesavento PA, Sullivan CS. 2015. Identification of a Polyomavirus microRNA Highly Expressed in Tumors. Virology 476: 43–53.

21. Bartel DP. 2004. MicroRNAs: genomics, biogenesis, mechanism, and function. Cell 116: 281–297.

22. Chen CJ, Cox JE, Kincaid RP, Martinez A, Sullivan CS. 2013. Divergent MicroRNA Targetomes of Closely Related Circulating Strains of a Polyomavirus. J Virol 87:11135–11147.

23. Broekema NM, Imperiale MJ. 2013. miRNA regulation of BK polyomavirus replication during early infection. Proc Natl Acad Sci U S A 110: 8200–8205.

24. Theiss JM, Günther T, Alawi M, Neumann F, Tessmer U, Fischer N, Grundhoff A. 2015. A Comprehensive Analysis of Replicating Merkel Cell Polyomavirus Genomes Delineates the Viral Transcription Program and Suggests a Role for mcv-miR-M1 in Episomal Persistence. PLoS Pathog 11:e1004974.

25. Zhang S, Sroller V, Zanwar P, Chen CJ, Halvorson SJ, Ajami NJ, Hecksel CW, Swain JL, Wong C, Sullivan CS, Butel JS. 2014. Viral MicroRNA Effects on Pathogenesis of Polyomavirus SV40 Infections in Syrian Golden Hamsters. PLoS Pathog 10: e1003912.

26. Bauman Y, Nachmani D, Vitenshtein A, Tsukerman P, Drayman N, Stern-Ginossar N, Lankry D, Gruda R, Mandelboim O. 2011. An identical miRNA of the human JC and BK polyoma viruses targets the stress-induced ligand ULBP3 to escape immune elimination. Cell Host Microbe 9:93–102.

27. Sung SK, Yim H, Andrews E, Benjamin TL. 2014. A Mouse Polyomavirus-encoded microRNA Targets the Cellular Apoptosis Pathway through Smad2 Inhibition. Virology 468–470: 57–62.

28. Gosert R, Rinaldo CH, Funk GA, Egli A, Ramos E, Drachenberg CB, Hirsch HH. 2008. Polyomavirus BK with rearranged noncoding control region emerge in vivo in renal transplant patients and increase viral replication and cytopathology. J Exp Med 205:841–852.

29. Chen CJ, Burke JM, Kincaid RP, Azarm KD, Mireles N, Butel JS, Sullivan CS. 2014. Naturally Arising Strains of Polyomaviruses with Severely Attenuated MicroRNA Expression. J Virol 88:12683–12693.

30. Swanson PA, Pack CD, Hadley A, Wang C-R, Stroynowski I, Jensen PE, Lukacher AE. 2008. An MHC class Ib-restricted CD8 T cell response confers antiviral immunity. J Exp Med 205:1647–1657.

31. Gattazzo F, Molon S, Morbidoni V, Braghetta P, Blaauw B, Urciuolo A, Bonaldo P. 2014. Cyclosporin A Promotes in vivo Myogenic Response in Collagen VI-Deficient Myopathic Mice. Front Aging Neurosci 244.

32. Kemball CC, Lee ED, Vezys V, Pearson TC, Larsen CP, Lukacher AE. 2005. Late priming and variability of epitope-specific CD8+ T cell responses during a persistent virus infection. J Immunol. 174:7950–60.

